# CTCF maintains pericentromere function and mitotic fidelity

**DOI:** 10.1101/2025.05.30.657091

**Authors:** Erin Walsh, Andrew D. Stephens

## Abstract

In mitosis the duplicated genome is aligned and accurately segregated between daughter nuclei. CTCF is a chromatin looping protein in interphase with an unknown role in mitosis. We previously published data showing that CTCF constitutive knockdown causes mitotic failure, but the mechanism remains unknown. To determine the role of CTCF in mitosis, we used a CRISPR CTCF auxin inducible degron cell line for rapid degradation. CTCF degradation for 3 days resulted in increased failure of mitosis and decreased circularity in post-mitotic nuclei. Upon CTCF degradation CENP-E is still recruited to the kinetochore and there is a low incidence of polar chromosomes which occur upon CENP-E inhibition. Instead, immunofluorescence imaging of mitotic spindles reveals that CTCF degradation causes increased intercentromere distances and a wider and more disorganized metaphase plate, a disruption of key functions of the pericentromere. These results are similar to partial loss of cohesin, an established component of the pericentromere. Thus, we reveal that CTCF is a key maintenance factor of pericentromere function, successful mitosis, and post-mitotic nuclear shape.

## Introduction

Mitosis is the stage of the cell cycle where duplicated tethered chromosomes are aligned and then segregated to opposite daughter cells. The mitotic spindle is made of microtubules emanating from opposite sides that must bind via the kinetochore to the centromere of sister chromatids, one centromere bound to each side, to cause biorientation. In metaphase, chromatin surrounding the centromere, termed the pericentromere, acts as a chromatin spring to properly resist microtubule spindle biorientation pulling forces to aid metaphase plate alignment (Lawrimore and Bloom, 2022; Andrade Ruiz *et al*., 2024). This alignment and tension are essential for accurate chromosome segregation in anaphase equally splitting the genome between the two daughter cells (Fonseca *et al*., 2019; Bunning and Gupta, 2023). Inaccurate chromosome segregation in mitosis can result in aneuploidy and cellular dysfunction that can promote human diseases like cancer (Levine and Holland 2018). While much is known about the mitotic spindle, kinetochore, and centromere, there are many proteins involved in this process that have yet to be determined.

CTCF is a zinc finger chromatin binding protein with a widely studied role in interphase but a relatively unknown role in mitosis. In interphase, CTCF controls chromatin looping behavior to regulate transcription (Hansen *et al*., 2017; Dehingia *et al*., 2022). CTCF and cohesin are known to interact to ensure proper looping of the chromatin (Gabriele *et al*., 2022). Initial studies have determined that CTCF has an essential role in mitosis (Wan *et al*., 2008; Xiao *et al*., 2015; Funk *et al*., 2022; Chiu *et al*., 2023; Watanabe *et al*., 2023). CTCF has been reported to localize to the pericentromere (Rubio *et al*., 2008; Xiao *et al*., 2015; Del Rosario *et al*., 2019; Miyata *et al*., 2021). The pericentromere is also a hotspot of chromatin looping by condensin and cohesin which aids sister chromatid biorientation and tensions sensing in metaphase (Lawrimore and Bloom, 2022). However, the exact role of CTCF in mitosis remains unclear.

Two core hypotheses remain untested for the mechanism underlying CTCF’s role in mitosis. One hypothesis is that CTCF is essential for recruiting CENP-E (Xiao *et al*., 2015) a chromo-kinesin that aids chromosome congression to the center of the mitotic spindle in metaphase (Yao *et al*., 1997). An alternative hypothesis is that CTCF, a known protein that aids chromatin looping, could aid pericentromere chromatin loop structure (Lawrimore and Bloom, 2019). Similar to its role in interphase, in mitosis CTCF might be interacting with cohesin at the pericentromere, where it is enriched. Destabilization of the pericentromere causes loss of tension sensing, increased intercentromere distance (Maresca and Salmon, 2009; Chiang *et al*., 2010), and abnormal separation in anaphase leading to lagging chromosome or anaphase bridges (Carvalhal *et al*., 2018).

To determine the role of CTCF in mitosis, we used a CTCF-mAID-Clover CRISPR cell line for rapid induction of CTCF degradation (Yesbolatova *et al*., 2020). Upon rapid degradation of CTCF, we time lapsed far-red labeled DNA to track mitotic failures, which increased significantly. We then measured nuclear shape via DNA fluorescence labeling to determine if CTCF knockdown caused changes in interphase nuclear shape properties. To determine the mechanism underlying CTCF’s role in mitosis, we conducted immunofluorescence of CENP-E to measure recruitment to the chromosomes and CENP-A to measure intercentromere stretch. Degradation of CTCF results in increased intercentromere distance and wider metaphase plate. Thus, we provide novel advancement in our understanding of the role of CTCF as a major contributor to pericentromere structure and function.

## Results and Discussion

### Multiple day degradation of CTCF increases mitotic errors

To rapidly degrade endogenous CTCF we used the previously generated CTCF-mAID-Clover HCT116 human cell line (Yesbolatova *et al*., 2020). Cells are treated with the auxin-inducible degradation system 2 (AID2) biochemical 5-Ph-IAA. Upon the addition of 5-Ph-IAA, >80% of CTCF-mAID-Clover intensity degrades within 2 hours (**Fig. 1, A and B**). This level of CTCF degradation remains constant over 3 days of 5-Ph-IAA treatment (**Fig. 1C**) and was confirmed via antibody immunofluorescence (**Supplemental Figure 1, A and B**). Thus, the AID2 system allows for rapid degradation of CTCF to determine short term effects on the scale of hours and days.

**Figure 1.**
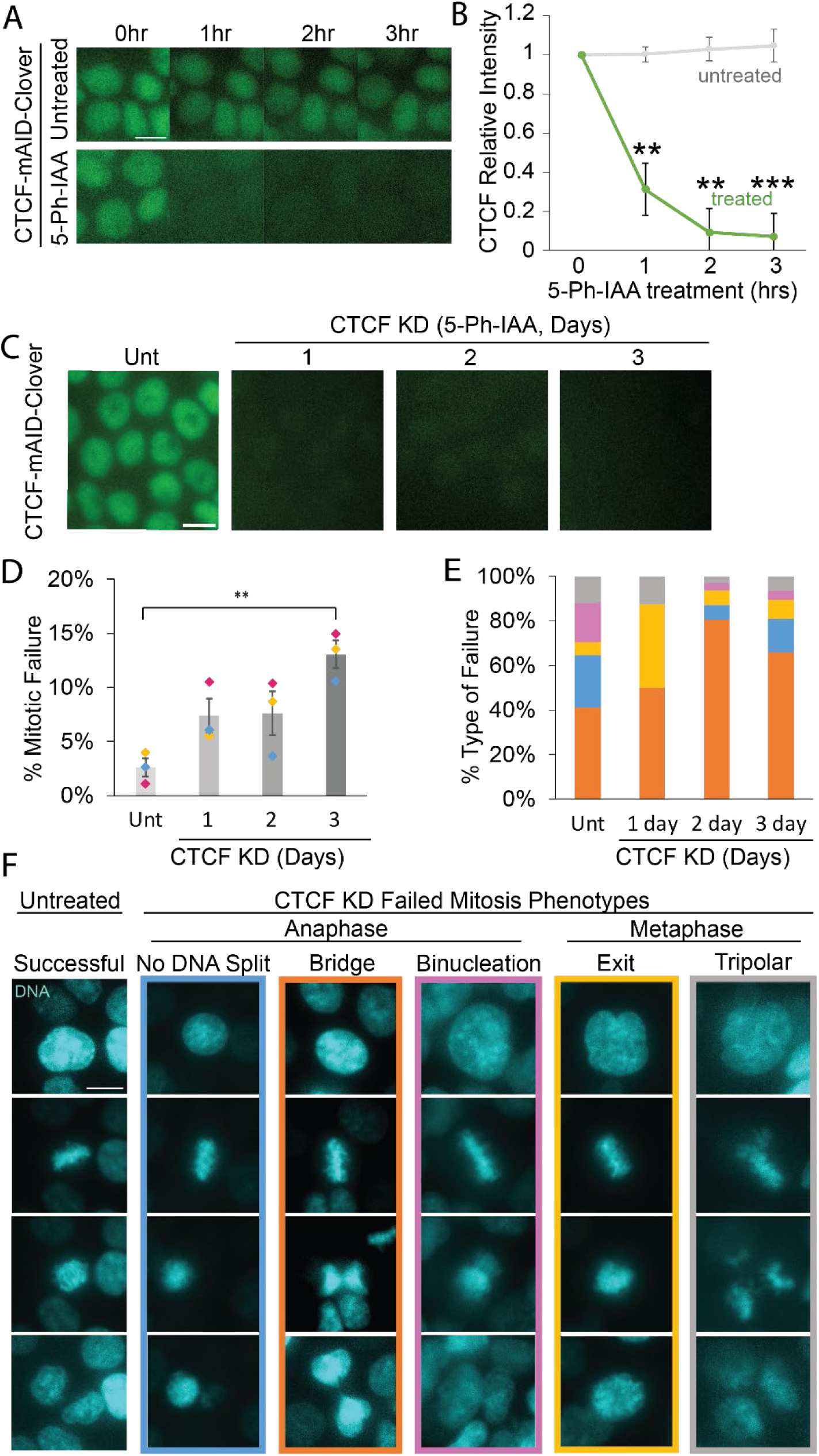
CTCF-mAID-Clover degradation causes increased mitotic failure rates mainly through anaphase failure. (A) Example images and (B) graph of CTCF-mAID-Clover relative fluorescence intensity at 0, 1, 2 and 3 hours in untreated (Unt) and 5-Ph-IAA treated. n >70 nuclei for each of 3 replicates. (C) Image examples CTCF-mAID-Clover fluorescence in untreated nuclei and nuclei treated with 5-Ph-IAA for 1, 2, and 3 days. (D) Graph of the rate of mitotic failure in untreated, 1-, 2-, and 3-day treatment with 10 µM 5-Ph-IAA. Data from three biological replicates shows failure levels of unt (8/199, 2/175, 6/223), 1 day (21/302), 2-day (14/161, 15/144, 5/135), and 3-day (14/103, 16/107, 16/151) treatment of 5-Ph-IAA for CTCF knockdown (CTCF KD). (E) Graph of the percentage of each type of mitotic failure in untreated, 1-, 2-, and 3-day CTCF KD. Colors represent the types of failures shown in panel F. The main phenotype, anaphase failure with DNA separation, are seen in untreated (7/17), 1 day (12/22), 2 day (25/31), and 3 day (31/47). (F) Example images of the 5 phenotypes of mitosis used to categorize mitotic failure or success. Images of nuclei stained with SPY-DNA and represented in cyan were taken from time lapse videos. Error bars represent standard error. Statistical tests are unpaired two-tailed Student’s t-test for panel B and E and ANOVA and Post Hoc Tukey tests for panel D. Significance is represented by *p < 0.05, **p < 0.01, and ***p < 0.001, and ns represents no statistical significance. Scale bar = 10 µm.

To determine if rapid loss of CTCF disrupts mitotic fidelity, we imaged cells stained with live cell DNA dye SPY650-DNA as they proceeded through mitosis. Cells were either untreated or treated with 5-Ph-IAA to degrade CTCF for 1 to 3 days. Time lapse imaging of SPY-DNA every 10 min for 16 hours revealed that untreated CTCF-mAID-Clover cells showed low levels of mitotic failure at 2.6 ± 0.8% (**Fig 1D**). Relative to untreated controls, mitotic errors increased slightly but insignificantly over 1 or 2 days of 5-Ph-IAA treatment to degrade CTCF (**Fig. 1D**).

CTCF degradation for 3 days resulted in a statistically significant increase in mitotic failure to 13.0 ± 1.2% (**Fig. 1D**). A 3-day 5-Ph-IAA treatment of the parent cell line, lacking the CRISPR modification of CTCF, did not change mitotic failure rates (untreated 1.6 ± 0.4% vs. parent 5-Ph-IAA 0.5 ± 0.5% mitotic failure, **Supplemental Figure 1C**). Thus, auxin induced degradation of CTCF results in a significant increase in mitotic failures after 3 days.

Mitotic failures were classified in groups dependent on the defining phenotype with the majority showing anaphase failures (**Fig. 1E**). The most prominent anaphase phenotype across conditions was an anaphase bridge between the daughter nuclei (orange, **Fig. 1E**). Less frequently, anaphase failed to split the DNA, resulting in all chromosomes going to one daughter nucleus and the other nucleus having none (blue, **Fig. 1E**). Another anaphase failure is binucleation in which daughter nuclei separated but remained in one cell, and thus the failure to separate two nuclei into two cells (magenta, **Fig. 1E**). Other minor CTCF knockdown phenotypes found were metaphase-based tripolar spindles and mitotic exit in which metaphase ends without attempting to separate the chromosomes or divide into two nuclei (gray and yellow respectively, **Fig. 1E**). Overall, CTCF degradation over multiple days caused increased mitotic failures but did not drastically change the distribution of types of mitotic failure. Furthermore, mitotic failures on average showed a delayed mitosis length relative to successful mitosis across conditions (**Supplemental Figure 1D**). Thus, the dominant failure upon CTCF degradation remained anaphase separation failures, largely consisting of anaphase bridges.

Discussion: CTCF degradation is known to have a delayed time interval. Studies of interphase looping showed that two days of CTCF degradation in ESC are required to affect 5C or Hi-C measurements of genome organization (Nora *et al*., 2017). The absence of effects upon rapid CTCF degradation suggests that it does not establish a key component of the mitotic spindle, like sister-sister cohesion or microtubule dynamics, both of which display immediate effects on mitosis (Salic *et al*., 2004; Cavazza *et al*., 2016; Zielinska *et al*., 2019; Yesbolatova *et al*., 2020; Chiu *et al*., 2023). Instead, CTCF likely plays a maintenance role that requires more than a single cell cycle to present significant changes to the mitotic spindle and its functionality. The data follows that loss of interphase looping occurring at 2 days CTCF degradation would lead to our observed increase, although insignificant, in mitotic errors at 2 days that become significant after 3 days of CTCF loss. Overall, loss of CTCF causes disruption of mitotic fidelity through anaphase failures agrees with many published reports that CFTC has a major role in mitotic segregation (Xiao *et al*., 2015; Chiu *et al*., 2023; Watanabe *et al*., 2023).

Anaphase bridges can lead to systemic genomic instability (Rodriguez-Muñoz *et al*., 2022). Concatenations that occur during DNA replication can lead to anaphase bridges. Usually, topoisomerase II cleaves the concatenated sisters aided by cohesin, condensin, and mitotic spindle microtubule tension to drive this reaction towards resolving sister chromatids separately (Finardi *et al*., 2020). Thus, CTCF could aid Topo II, cohesin, and condensin in mitosis to resolve sister chromatids under tension. Disruption of the kinetochore causing the inability to sense tension through kinetochore stretching has been shown to cause anaphase bridges as well (Uchida *et al*., 2021). One possibility is that loss of CTCF disrupts the pericentromere chromatin spring causing decreased tension sensing at the kinetochore. Partial loss of cohesin during mitosis, an essential component of the pericentromere chromatin spring, has been shown to produce anaphase bridges, lagging chromosomes, and extended mitosis length (Carvalhal *et al*., 2018). Our data of mitotic failure and phenotype upon loss of CTCF support a possible role in sister chromatid decatenation and/or aiding cohesin in pericentromere structure to resist tension.

### CTCF long-term degradation leads to abnormally shaped nuclei from post-mitotic nuclear reformation

We reasoned that mitotic failures or even slight disruptions in mitosis could manifest as post-mitosis nuclear reformation abnormal nuclear shape, a hallmark of human disease that can cause dysfunction (Stephens *et al*., 2019; Kalukula *et al*., 2022). To accomplish this, we measured post-mitosis nuclear reformation shape via nuclear circularity 1.5 hours after anaphase onset. A measurement of 1.0 nuclear circularity is a perfect circle with all concave edges while convex edges decrease this value **(Fig. 2A)**. Nuclear circularity average measurements showed significant changes between untreated and 3-day CTCF knockdown **(Fig. 2B)**. Individual nuclei data show a growing tail into lower nuclear circularity, or more abnormally shaped nuclei. Using the threshold of abnormal nuclear shape < 0.85 circularity, we show a drastic increase from untreated with 4.5 ± 3.0% to CTCF knockdown 31.9 ± 2.6% abnormally shaped nuclei post-mitosis nuclear reformation (**Fig. 2C**). Nuclear size distributions changed slightly but insignificantly over this time interval for degradation (**Supplemental Figure 1E**). Thus, taken together CTCF loss disrupts both mitosis and post-mitosis nuclear shape in interphase.

**Figure 2.**
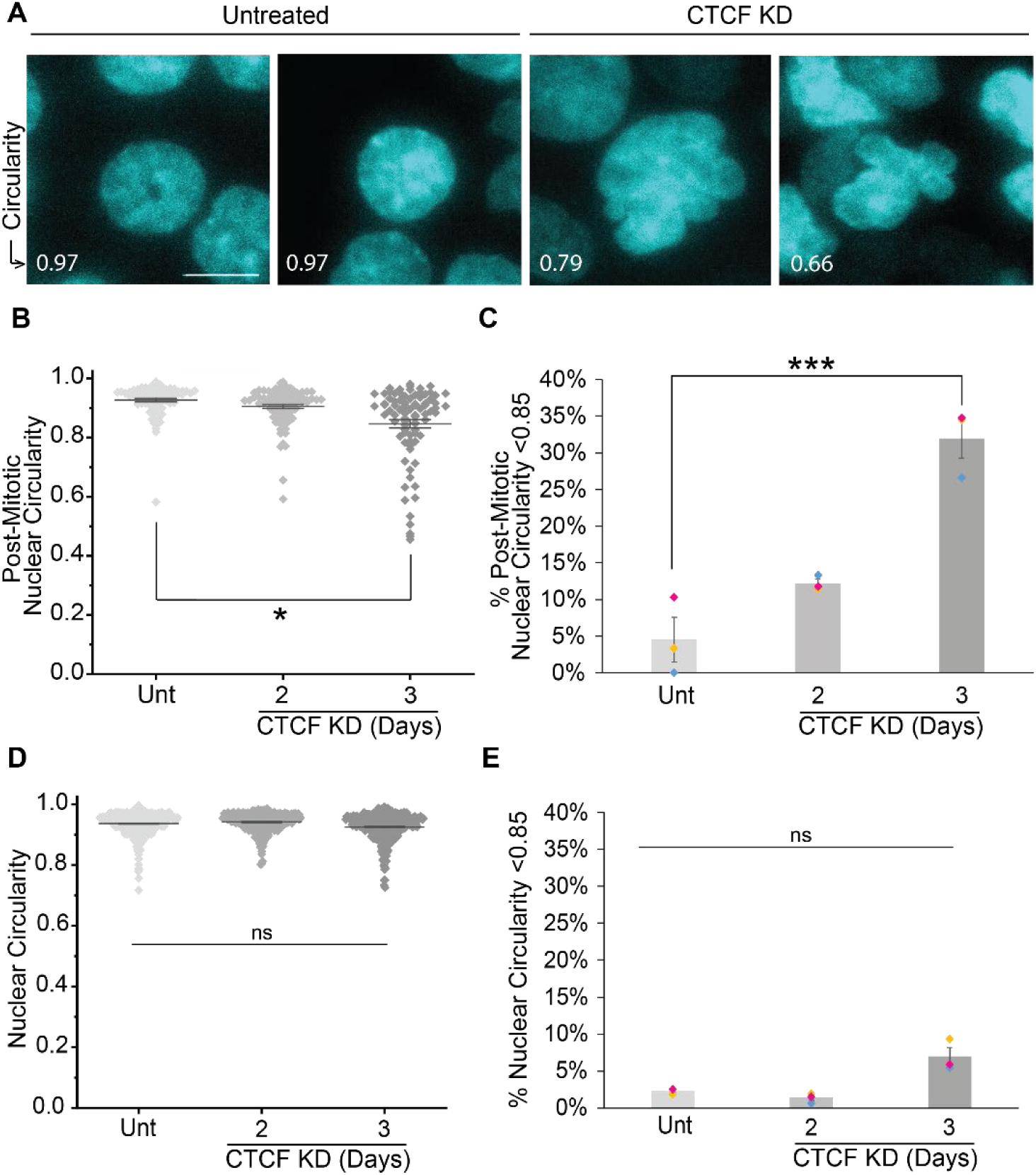
CTCF knockdown after 3 days causes decreased nuclear circularity post- mitosis. (A) Example images highlighting the circularity differences between untreated and 3- day 5-Ph-IAA treatment for CTCF knockdown (CTCF KD). Nuclei are marked with SPY-DNA (cyan) and circularity measurement is shown in the bottom left corner. Scale bar is 10 µm. (B) Graph of post-mitotic circularity measurements 1.5 hours after anaphase onset of untreated (Unt) and 2 or 3 days CTCF knockdown. (C) Graph of the percentage of nuclei with post-mitotic nuclear circularity < 0.85. The data from B and C are from three replicates each with n > 16 from the 16-hour time lapse. Post-mitotic nuclear circularity < 0.85 in untreated (5/89), 2-day (11/90), and 3-day (27/82) CTCF KD. (D,E) Graphs of (D) nuclear circularity measurements and (E) percentage of nuclei with circularity < 0.85 to denote abnormal for untreated and 2- or 3-day CTCF KD. Data are measured from total nuclei population at one time point for three replicates each with n > 350. Number of nuclei below the threshold are untreated (11/461), 2-day (5/379), 3-day (28/423) CTCF KD. Statistical tests for panel B were Kruskal Wallis and Dunns tests. Statistical tests for C-E were ANOVAs followed by Post Hoc Tukey tests. Error bars represent standard error. Significance is represented by *p < 0.05, **p < 0.01, and ***p < 0.001, and ns represents no statistical significance.

To determine if CTCF also causes abnormal nuclear shape changes in interphase, we imaged the population of interphase nuclei. Interphase nuclei populations showed no change in average nuclear circularity (**Fig. 2D**). Furthermore, we measured the percentage of nuclei with a circularity < 0.85 to address the possibility of an increase in the subpopulation of abnormally shaped nuclei. Untreated compared to 3-day CTCF knockdown resulted in no significant changes in nuclei with a circularity < 0.85 (**Fig. 2E**). Thus, CTCF loss has no significant effect on population nuclear shape but does cause a significant increase in mitotic failure and post-mitosis abnormal nuclear shape.

Discussion: Mitotic failures are well known to result in abnormal nuclear shape (Gisselsson *et al*., 2001). Thus, CTCF’s role in mitosis has downstream implications for the cell in interphase when the miotic chromosome decondense and reforms into the nucleus. Abnormal nuclear shape has been shown to result from disruption of chromosome alignment in metaphase that persists into ana/telophase and ultimately nuclear reformation which can lead to dysfunction (Fonseca *et al*., 2019). Abnormal nuclear morphology is a hallmark of human disease. Recent findings have revealed that abnormal nuclear shape via interphase-based nuclear blebs is not only a hallmark but can be a driver of disease progression and overall cellular dysfunction (Srivastava *et al*., 2021; Pho *et al*., 2023; Chu *et al*., 2025). Thus, abnormal shape via mitosis could cause similar dysfunctions. Finally, it is well known that CTCF disrupts chromosome organization measures by Hi-C (Hansen *et al*., 2017; Pugacheva *et al*., 2020; Gabriele *et al*., 2022; Banigan *et al*., 2023). It is possible that perturbed nuclear shape upon reformation is a key disruptor of chromosome organization that is being missed by shorter CTCF degradation experiments of which there are many published. Overall, CTCFs role in accurate mitotic segregation might extend into the interphase nucleus to determine function.

### CTCF is not required for CENP-E recruitment or function

Previous publications have shown that CTCF has a role in mitotic fidelity, but a mechanism remains unclear. One hypothesis for the mechanism behind CTCF’s role is localizing CENP-E to the centromere, supported by one publication (Xiao *et al*., 2015). Recruitment of CENP-E can be determined via immunofluorescence of CENP-E during mitosis imaged via spinning disk confocal microscopy (**Fig. 3A**). CENP-E immunofluorescence in the metaphase plate showed no significant change from untreated to 3-day CTCF degradation via 5-Ph-IAA (**Fig. 3, A and B**). However, the inhibition of CENP-E via GSK 923295 for 1 day resulted in a drastic and significant 60% loss of CENP-E intensity in the metaphase plate chromosomes (**Fig. 3B**). Thus, our data does not support a role for CTCF in controlling CENP-E levels at the kinetochore in metaphase.

**Figure 3.**
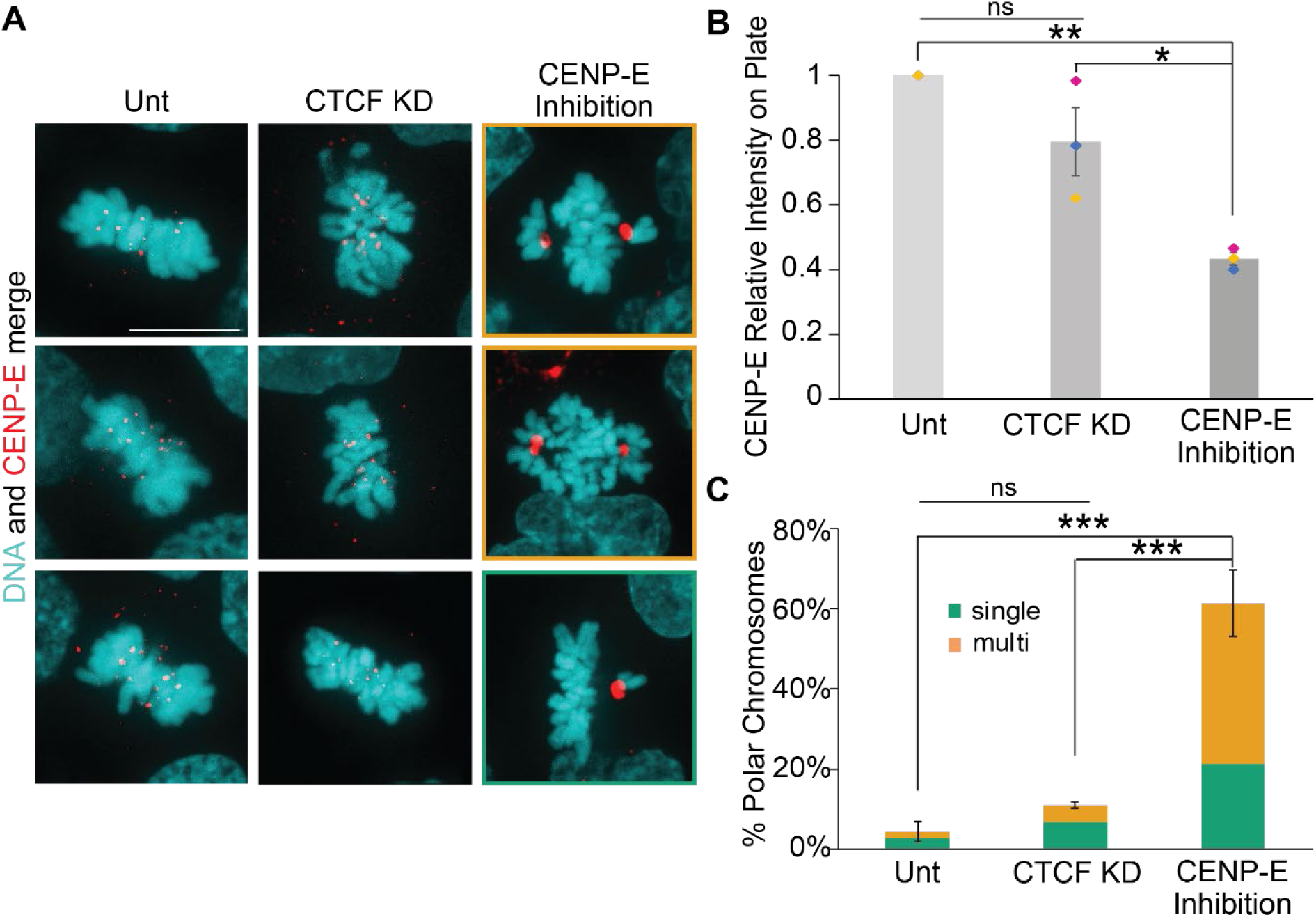
CENP-E intensity and chromosome congression remain intact upon CTCF knockdown. (A) Metaphase example images of CENP-E immunofluorescence (red) and DNA stained with Hoechst (cyan) in untreated (Unt), CTCF KD via 3-day 5-Ph-IAA treatment, and CENP-E inhibitor GSK-923295 treatment 10nm for 1 day. Scale bar is 10 µm. (B) Graph of CENP-E relative intensity on metaphase plate chromosomes in untreated, CTCF KD, and GSK- 923295. Data are from 3 biological replicates with n>12 for each replicate. (C) Graph of the percentage of polar chromosomes and categorizations of single (green) or multiple (orange) polar chromosome pairs present in untreated, CTCF KD, and GSK-923295. Data for polar chromosomes represents three replicates with n>15 for each replicate. Polar chromosomes are present in untreated (3/69), 3-day CTCF KD (5/27), GSK-923295 (63/98). Statistical tests for panel B and C ANOVAs followed by Post Hoc Tukey tests. Error bars represent standard error. Significance is represented by *p < 0.05, **p < 0.01, and ***p < 0.001, and ns represents no statistical significance.

CENP-E is a chromokinesin that loads at the spindle pole, binds to chromosomes, and walks them down the spindle microtubules towards the center of the spindle allowing chromosomes to congress for biorientation and metaphase plate alignment. Failure of this process results in polar chromosomes, defined as chromosomes that fail to congress to the metaphase plate and remain near the spindle pole (Barisic *et al*., 2014). The phenotypic changes in mitosis due to disruption of CENP-E can be determined by treatment with a CENP-E inhibitor. Treatment with CENP-E inhibitor GSK 923295 for 1 day resulted in polar chromosomes in 62% of cells in metaphase, with the majority of which had more than one sister chromatid pair at the spindle poles (**Fig. 3, A and C**). CENP-E accumulation at polar chromosomes aids identification and shows lack of function. This data agrees with many publications showing CENP-E knockdown or inhibition results in polar chromosomes (Bennett *et al*., 2015). CTCF knockdown for 3 days resulted in a modest but insignificant increase in the presence of polar chromosomes in metaphase from 4% untreated to 13% in CTCF KD with the majority showing only a single chromatid pair at a spindle pole (**Fig. 3C**). Thus overall, loss of CTCF presents an insignificant change while inhibition of CENP-E via GSK 923295 leads to drastic increase in polar chromosomes in the majority of metaphase spindles. Taken together, the data does not support a role for CTCF in CENP-E recruitment or function.

Discussion: The disagreement between our data showing no dependence and previous reports of CENP-E dependence on CTCF might be due to the difference in CTCF loss time. Our data via 3-day CTCF knockdown supports that CENP-E does not rely on CTCF for recruitment or function. Our novel data looks at the first few cell cycles after loss of CTCF via rapid degradation via auxin-inducible degradation. The other publications showing the importance of CTCF in mitosis are the result of longer-term loss of CTCF (Xiao *et al*., 2015; Chiu *et al*., 2023; Watanabe *et al*., 2023). Thus, it is possible the loss of CTCF over more than 2-3 cell cycles causes a set of secondary effects to disrupt other centromere and kinetochore structures and/or functions. Our data is particularly novel in that with a few days CTCF knockdown, we find that the primary role of CTCF is not CENP-E recruitment but likely a different function in the pericentromere where it is enriched in mitosis.

### CTCF is essential to pericentromeric structure and metaphase plate alignment

An alternative hypothesis is that CTCF aids cohesin in maintaining the pericentromere chromatin spring. Pericentromeric structure can be directly measured by intercentromere distance imaged via spinning disk confocal microscopy. Using Anti-Centromere Antibody (ACA) we used immunofluorescence to label and measure the distance between sister centromeres (**Fig. 4A**). In untreated cells the intercentromere distance measured between bioriented sister chromatid centromeres in metaphase was 1.08 ± 0.01 µm. CTCF knockdown for 3 days resulted in significant increase in intercentromere distance to 1.26 ± 0.04 µm (**Fig. 4B**). This data supports the hypothesis that CTCF has a role in maintaining pericentromere chromatin structure as a spring to resist bioriented microtubule pulling forces.

**Figure 4.**
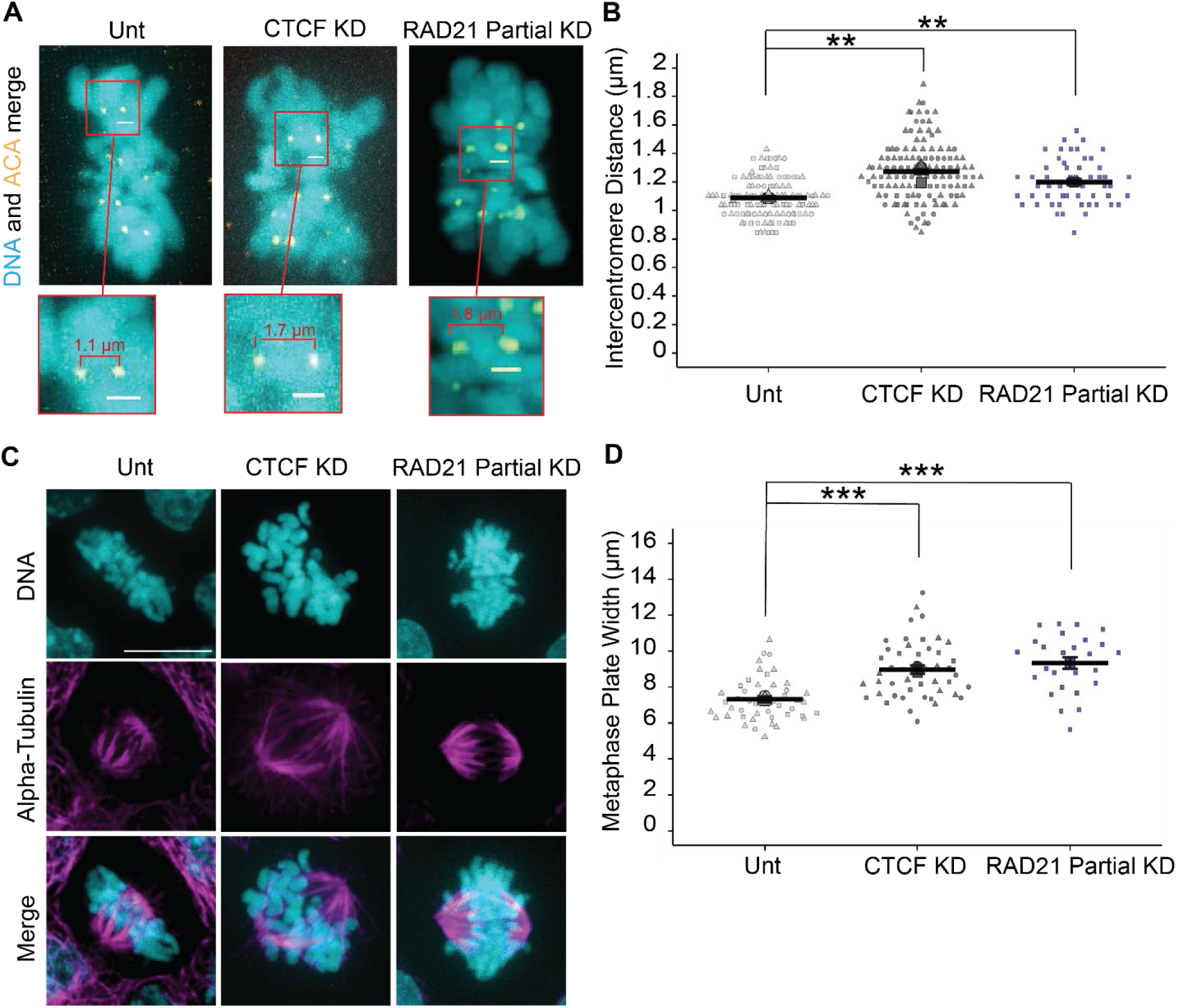
Intercentromere distance and metaphase plate width increase with CTCF knockdown and RAD21 partial knockdown. (A) Example images of sister centromere pairs and (B) graph of intercentromere distances in metaphase plates in untreated (Unt), 3-day 5-Ph-IAA (CTCF KD), and RAD21 partial knockdown via 6 hours 5-Ph-IAA 357µM treatment. DNA is stained with Hoechst (cyan) and the centromere is marked by ACA immunofluorescence (yellow). Scale bar is 1 µm. Data for unt and CTCF KD are from three biological replicates with n>10 for each replicate. Data for RAD21 partial KD are from one biological replicate with n>26. RAD21 partial knockdown data is provided in Supplemental Figure 1F. (C) Example images of metaphase plates in which DNA is stained with Hoechst (cyan), and immunofluorescence marks alpha tubulin (magenta) in untreated, CTCF KD, RAD21 partial KD. Scale bar is 10 µm. (D) Graph of metaphase plate width. Data for Unt and CTCF KD are from three biological replicates with n>15 and RAD21 partial KD are from one replicate n= 16. Statistical tests run were unpaired two-tailed Student’s t-tests between untreated and each KD. Error bars represent standard error. Significance is represented by *p < 0.05, **p < 0.01, and ***p < 0.001, and ns represents no statistical significance.

The pericentromere spring aids in both tension sensing and alignment of chromosomes in metaphase. We measured the metaphase plate width using spinning disk confocal imaging of DNA labeled via Hochest. Untreated metaphase plate width visually appears tight, and measured 7.33 ± 0.07 µm (**Fig. 4, C and D**). CTCF knockdown for 3 days resulted in visually disrupted metaphase plates which increased significantly in width to 8.96 ± 0.08 µm relative to untreated (**Fig. 4, C and D**). Overall, loss of CTCF disrupts chromosome organization on the metaphase plate and increases intercentromere distance, two roles dedicated to the pericentromeric chromatin spring which is also maintained by cohesin.

Partial loss of cohesin has been shown to have a role in determining the pericentromere chromatin spring structure and function (Carvalhal *et al*., 2018). To recapitulate those findings and compare to CTCF KD, we used previously generated CRISPR cell line RAD21-mAID-Clover HCT116 cells (Yesbolatova *et al*., 2020) and treated with 357 µM of 5-Ph-IAA for 6 hours to partially degrade cohesion (**Supplemental Figure 1F**). Upon RAD21 partial knockdown, we then measured intercentromere distance and metaphase plate width. Partial cohesin degradation resulted in a significant increase relative to untreated in both intercentromere distance to 1.19 ± 0.02 µm and metaphase plate width to 9.33 ± 0.31 µm, similar to CTCF degradation (**Fig. 4, B and D**). In summary, knockdown of CTCF and partial knockdown of cohesin result in increased intercentromere distance and metaphase plate width.

DISCUSSION: CTCF is known to be enriched in the pericentromere (Rubio *et al*., 2008; Xiao *et al*., 2015; Del Rosario *et al*., 2019; Miyata *et al*., 2021). However, the role of CTCF in the pericentromere remained unknown until now. Our data provides strong support for CTCF’s role in maintaining pericentromere structure and function through intercentromere distance and metaphase plate alignment. Disruption of the kinetochore has been shown to decrease interkinetochore/centromere distance (Maresca and Salmon, 2009). This outcome is likely due to the inability to attach and biorient sister chromatids to pull them in opposite directions to result in less stretching or distance. Considering this, our data show that CTCF is not essential for kinetochore attachment. Instead, attached and bioriented centromeres cannot resist this tension due to a weakened pericentromere chromatin spring. It has been established in yeast that the pericentromere spring is composed of cohesin and condensin loops on loops (Yeh *et al*., 2008; Stephens *et al*., 2011, 2013b, 2013a; Lawrimore *et al*., 2018). Partial loss of cohesin, where sister cohesion is maintained, results in increased inter-kinetochore/centromere distance and mitotic errors (Carvalhal *et al*., 2018; Zielinska *et al*., 2019). Thus, loss of CTCF recapitulates partial cohesin loss which we have recapitulated in our data. We provide novel data that the role of CTCF in mitosis is similar to its role in interphase, to aid cohesin based looping, though in mitosis this role extends to maintain the pericentromere chromatin spring. This data overall suggests that CTCF maintains the pericentromere function via sister centromere tension and metaphase plate alignment to aid proper mitotic fidelity and normal nuclear shape post-mitosis.

## Materials and Methods

### Cell culture

HCT116 WT and HCT116 CTCF- mAID-Clover and RAD21-mAID-Clover cells were grown in McCoys 5A media (Thermofisher Scientific, 16600-082) that was supplemented with 10% fetal bovine serum (FBS; HyClone) and 1% penicillin/streptomycin (Corning). The cells were allowed to grow to 70% confluency on 60mm dishes before being passaged every 2-3 days.

### Drug Treatment

The drug of interest for this paper is 5-Ph-IAA (Sigma Aldrich, SML3574), the inducer of the AID system. For the CTCF-mAID-Clover cells, cells were plated and allowed to grow on the dish for 24 hours before being treated with 10uM of 5-Ph-IAA for 1-3 days depending on the condition. Media was aspirated, replaced, and re treated every 24 hours to ensure proper functioning of the 5-Ph-IAA. For the RAD21-mAID-Clover cells, cells were plated and allowed to grow on the dish for 24 hours before being treated with 357µM of 5-Ph-IAA for 6hrs.

GSK-925295 (APExBIO, a3450) was used to represent CENP-E inhibition phenotypes. Cells were plated, allowed to grow for 24hrs, and then treated with 10 nM of GSK for 1 day before imaging.

### Time-lapse imaging and analysis

Images were acquired with Nikon Elements software on a Nikon Instruments Ti2-E microscope, Orca Fusion Gen III camera, Lumencor Aura III light engine, TMC CLeanBench air table, with 40x air objective (N.A 0.75, W.D. 0.66, MRH00401). Live cell time lapse imaging was possible using Nikon Perfect Focus System and Okolab heat 37°C, humidity, and 5% CO2 stage top incubator (H301).

For rapid degradation imaging, cells were plated in four well imaging dishes (Cellvis, D35C4-20- 1.5-N), and allowed to grow for one day. The cells were then treated with SPY650-DNA (Thermofisher Scientific, NC2096299) and 5-Ph-IAA, and imaging began after 45 minutes.

Images were taken in 7-minute intervals for 3 hours with 5 Fields of view per condition. Cells were imaged using the Cy5 channel to visualize the DNA, and FITC to visualize the CTCF-mAID-Clover. ROIs were drawn around nuclei according to the DNA staining, and the FITC intensities were taken throughout the time lapse. Data was transferred to excel and background subtracted. The percent intensity at the various time points was compared to the intensity at T=0.

For mitotic failure imaging, cells were plated in four well imaging dishes and allowed to grow for one day before being treated with 5-Ph-IAA for various days. Cells were then treated with SPY650-DNA and imaging commenced after 45 minutes. Images were taken in 10-minute intervals for 16 hours with 7 fields of view per condition and a 3-step z-stack of 2.5 µm steps to cover 5 µm. Cells were imaged using the Cy5 channel to visualize DNA as well as the consolidation of DNA associated with the steps of mitosis.

Analysis of mitotic duration was calculated from the condensation of the DNA to the abscission of the two nuclei, or in the case of failed mitosis the eventual expansion of the DNA. The number of frames from beginning to end was multiplied by the length of the frame (10 minutes), and this data was noted in Excel (Microsoft).

Mitotic failure was categorized by the five categories and noted in Excel, and the activity of the nuclei were confirmed to match to each category by observing their motion for one hour post mitosis.

Area and circularity for each condition was calculated at the beginning (t=0) of each time lapse, when the treatment was exactly 48 or 72hrs (corresponding to 2, and 3 days). To do this, a Bezier ROI was drawn around each nucleus that did not overlap any other nucleus, converted to binary, and then pasted into excel where the data from all 3 replicates were compiled. For post mitotic circularity, the nuclei were allowed to exit mitosis, and then 10 frames (100 minutes) post mitosis an ROI was hand drawn over any resulting individual daughter nuclei. The ROI was converted to binary, and the data was pasted into excel.

### Immunofluorescence

Cells were plated in 8 well imaging dishes (Cellvis, C8-1.5H-N) and allowed to grow for one day before treatment with 5-Ph-IAA for multiple days. Cells were then fixed in 4% paraformaldehyde (Electron Microscopy Sciences), washed with Phosphate-Bufferd Saline (PBS; Corning) three times for 5 minutes each. Cells were permeabilized with 0.1% Triton X-100 (Promega) for 15 minutes, then with 0.06% Tween (PBS-T) for 5 minutes, and then three washing steps with PBS were completed. Cells were then blocked in 2% Bovine Serum Albumin (BSA; Thermofisher Scientific) in PBS for one hour at room temperature.

Primary antibody dilutions were created using BSA in PBS. The primary antibodies used were CTCF Rabbit (2899, Cell Signaling Technologies) 1:100, CENP-E mouse (39619, Active Motif) 1:1000, ACA human (HCT-0100 Immunovision) 1:4000, CST) 1:1000. Primaries were added and allowed to sit overnight at 4 degrees. Secondary antibody dilutions were created with BSA in PBS and treated with concentrations of 1:1000 for 1 hour on a shaker at room temperature. The secondary antibodies used were goat anti-human 555 (A21433 Invitrogen), goat anti-mouse 647 (4410, Cell Signaling Technologies), goat anti-rabbit 647 (4414, Cell Signaling Technologies). Cells were then washed 3 times with PBS.

The cells were then stained with 1:10,000 concentration of Hoechst 33342 (H3570, Thermofisher Scientific) for 10 minutes before 3 more PBS washing steps. The cells were then mounted using 30uL Gold Antifade per well (P36930, Thermofisher Scientific).

### Immunofluorescence imaging and analysis

As previously described in (Pho *et al*., 2023), spinning disk confocal images were acquired with Nikon Elements software on a Nikon Instruments Ti2-E microscope with Crest V3 Spinning Disk Confocal, Orca Fusion Gen III camera, Lumencor Celesta light engine, TMC CleanBench air table, with Plan Apochromat Lambda 100× Oil Immersion Objective Lens (N.A. 1.45, W.D. 0.13 mm, F.O.V. 25 mm, MRD71970).

The metaphase plate stained by Hoechst was visualized in DAPI and imaged with a 100x oil objective. The width of the metaphase plate was calculated by creating a maximum projection image in Z with DNA, and then measuring the distance between the edges of DNA using the length tool in annotations and measurements in NIS. This measurement includes the metaphase tail that is present in most plates and also includes any chromosomes that were not successfully connected to the metaphase plate during the transition from prophase to metaphase. This ROI was converted to binary, where its width measurement was taken and compiled into excel.

The centromeres were visualized in TRITC by staining of Anti-Centromere Antibody (ACA). They were imaged with a z stack of 0.3um steps for 5um, and imaged with a 100xoil objective with settings of 60% 300ms 12-bit. The intercentromere distance was measured by cropping an image of a metaphase plate, drawing an ROI where only background fluorescence was present in both DAPI and TRITC. This ROI was set as a background ROI and then used to background subtract the image. A line scan was then drawn through two foci that were in the same plane, on the metaphase plate, and had a visible chromosome between them. The distance between the peak intensities of the two foci was calculated and pasted in excel as the intercentromeric distance for that one pair. This was completed 1-3 times per metaphase plate

CENP-E was visualized in CY5 and imaged with a 100x oil objective with settings of 99% 350ms 12-bit-sensitive. The metaphase plate was cropped in NIS elements, and the background signal was subtracted by drawing an ROI, setting it as a background ROI, and subtracting that signal from the image. A max project in Z was created with DNA and CENP-E, and then a bounding box was drawn around the main metaphase plate (excluding polar chromosomes and tails). The intensity of CENP-E in each plate was compiled and analyzed in excel.

### Statistics

After data was compiled, Shapiro-Wilks normality tests as well as Levene Tests for homogeneity of variance were conducted on RStudio. If the data sets did not pass this test, a Kruskal Wallis test was run followed by a Dunn’s test. If the data sets passed these normality tests with a p value > 0.05, the data sets were tested for statistical significance using an ANOVA test. If the resulting p value was >0.05, then the data was not statistically significant. If the resulting p value was < 0.05, then the data was statistically significant, and a Post Hoc Tukey Test was run to test for significance between each group. A group was considered significantly different from another if the p value was < 0.05. If the data did not have 3 replicates, a students ttest was run instead. The statistical tests run on each data set are noted in each figure legend. To summarize, ANOVA and Post Hoc Tukey tests were run on Fig 1D, Fig 2C-E, FIG 3B and 3C, Fig 4D, and Supplemental Fig 1E. Kruskal Wallis and Dunns tests were run on Fig 2B, and Fig 4B. Students t-tests were run on Supplemental Fig 1B-D and 1F.

## Acknowledgements

We would like to thank Masato T. Kanemaki for kindly sharing the CTCF-mAID-Clover and RAD21-mAID-Clover cell lines. We would like to thank Thomas Maresca and Thomas Laskarzewski for helpful and insightful discussions. We would like to thank Kelsey Prince for feedback and support as a great lab mate.

## Data availability

All figure data can be found posted in the public repository FigShare: https://doi.org/10.6084/m9.figshare.29197922.v1

