## Supplemental Figure 1 for "CTCF maintains pericentromere function and mitotic fidelity"

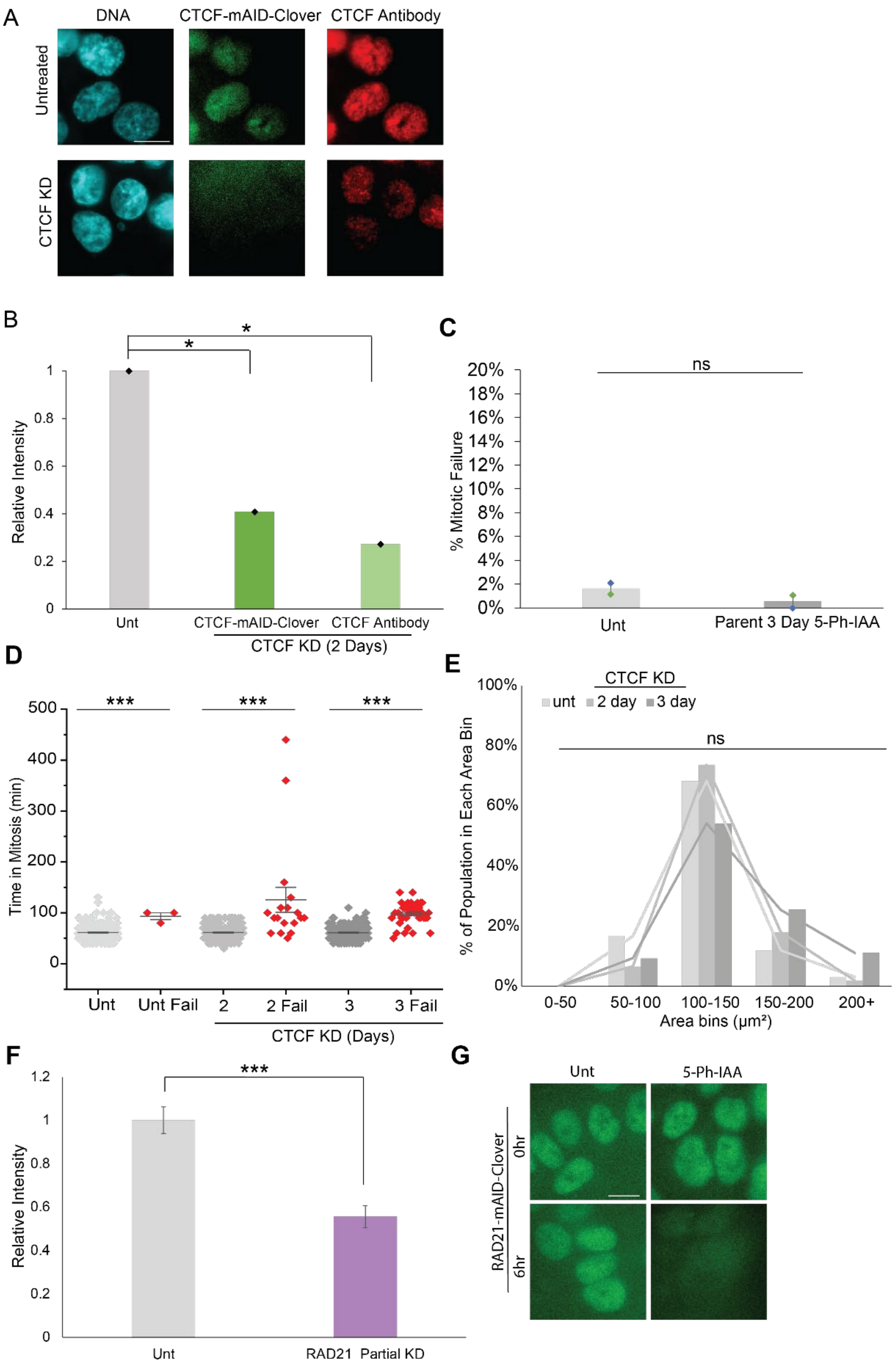

A
DNA
CTCF-mAID-Clover
CTCF Antibody
Untreated
CTCF KD
B
Relative Intensity
Unt
CTCF-mAID-Clover
CTCF Antibody
CTCF KD (2 Days)
C
% Mitotic Failure
ns
Unt
Parent 3 Day 5-Ph-IAA
D
Time in Mitosis (min)
Unt
Unt Fail
2
2 Fail
3
3 Fail
CTCF KD (Days)
E
% of Population in Each Area Bin
CTCF KD
unt
2 day
3 day
ns
0-50
50-100
100-150
150-200
200+
Area bins (µm²)
F
Relative Intensity
Unt
RAD21 Partial KD
G
Unt
5-Ph-IAA
RAD21-mAID-Clover
0hr
6hr

**Supplemental Figure 1. CTCF-mAID-Clover degradation controls, mitotic length, and nuclear size.** (A) Example images and (B) graph of relative intensity levels via CTCF-mAID-clover and CTCF antibody immunofluorescence after 2 days of 5-Ph-IAA treatment. Data from one biological replicate with n>20. (C) Data from two biological replicates show the number of failed mitotic events in untreated (1/85, 2/93) and 3-day CTCF KD by 5-Ph-IAA treatment (1/93, 2/95). (D) Data from three biological replicates show the average time a nucleus spends in mitosis during successful events (gray) and failed mitotic events (red). (E) Histogram of nuclear size bins as a total percentage of untreated or 2-3 day CTCF KD nuclei. Data represents 3 biological replicates with n>100 for reach replicate. (F) Graph representing the relative intensities of untreated nuclei and nuclei treated with 5-Ph-IAA for 6 hours to partially degrade RAD21. Data is from one biological replicate with n>26. (G) Example images of untreated nuclei and nuclei treated for 6 hours with 5-Ph-IAA to partially degrade RAD21. Scale bar is 10 μm. Statistical tests run in B-D and F were unpaired two-tailed Student's t-tests. Statistical tests run for E were ANOVAs followed by Post Hoc Tukey tests. Error bars represent standard error. Significance is represented by *p < 0.05, **p < 0.01, and ***p < 0.001, and ns represents no statistical significance. Scale bar is 10 μm.
